# Cochaperones convey the energy of ATP hydrolysis for directional action of Hsp90

**DOI:** 10.1101/2023.06.16.545234

**Authors:** Leonie Vollmar, Julia Schimpf, Bianca Hermann, Thorsten Hugel

**Affiliations:** Institute of Physical Chemistry, University of Freiburg, Germany; Signalling Research Centers BIOSS and CIBSS, University of Freiburg, Germany; Spemann Graduate School of Biology and Medicine (SGBM), University of Freiburg

## Abstract

The molecular chaperone and heat shock protein Hsp90 is part of many protein complexes in eukaryotic cells. Together with its cochaperones, Hsp90 is responsible for the maturation of hundreds of clients. Although having been investigated for decades, it still is largely unknown which components are necessary for a functional complex and how the energy of ATP hydrolysis is used to enable cyclic operation. Here we use single-molecule FRET to show how cochaperones introduce directionality into Hsp90’s conformational changes during its interaction with the client kinase Ste11. Most interestingly, three cochaperones are needed to couple ATP turnover to these conformational changes. All three are therefore essential for a functional cyclic operation, which requires coupling to an energy source. Finally, our findings show how the formation of sub-complexes in equilibrium followed by a directed selection of the functional complex can be the most energy efficient pathway for kinase maturation.

## Introduction

Being one of the most abundant proteins in the cytoplasm (Borkovich et al. 1989), the molecular chaperone and heat shock protein Hsp90 collaborates with more than 20 cochaperones to help other proteins (clients) acquire their active conformation (Taipale et al. 2010; Morán Luengo et al. 2019). The homodimeric Hsp90 is structurally well-equipped for assisting numerous clients. Each Hsp90-monomer consists of three domains: The N-terminal domain (NTD) contains the ATP-binding site, the middle domain (MD), which is connected to the NTD by a charged linker sequence, is important for ATP-hydrolysis (Hoter et al. 2018), and the C-terminal domain (CTD), which is the dimerization interface of highest affinity (Hoter et al. 2018), contains a C-terminal MEEVD motif (Scheufler et al. 2000). Due to its structural flexibility, the Hsp90-dimer can undergo large conformational changes ranging from an open, V-shaped structure to a tightly closed state (Ali et al. 2006; Krukenberg et al. 2011; Mickler et al. 2009), where also the middle and N-terminal domains dimerise.

During the last years, several models were suggested in which Hsp90’s client processing as well as related structural changes are linked to an ATP hydrolysis cycle. However, it is still unclear which components of the Hsp90 machinery are necessary for an efficient use of the energy from ATP hydrolysis. In fact, one of the long standing questions in the field is how ATP hydrolysis is utilized in the Hsp90 machinery at all. Similar questions have been answered for motor proteins (Phillips et al. 2013), GTPases (Singh et al. 2023) or DNA gyrase (Basu et al. 2018) but not for chaperones, where a clear directional coordinate is missing. Even for specific complexes like the Hsp90-cochaperone-kinase machinery, the whole dynamic picture remains enigmatic and its depiction highly suggestive. Several publications propose, for example, that the complex closes upon ATP-binding (Verba and Agard 2017) and then rearranges to a second, more compact closed structure (Wolf et al. 2021). Upon ATP-hydrolysis, the complex opens again and releases the maturated kinase as well as the cochaperone, leaving Hsp90 ready for yet another cycle (Verba and Agard 2017; Eckl et al. 2015; Keramisanou et al. 2022; Schopf et al. 2017). Alternatively, it was shown that Hsp90 does not close upon ATP binding itself (Lopez et al. 2021), but instead (Reidy et al. 2023) proposed recently that its conformational changes are triggered by the proper positioning of ATP’s gamma phosphate. Furthermore, it was proposed that the cycle might also be linked to dephosphorylation of a kinase (Jaime-Garza et al. 2023). Such cycles are a reasonable assumption but have not yet been shown experimentally. They imply directionality, i.e. entropy production at constant turnover, which can only be proven with single-molecule experiments as detailed in the following.

To better understand the concept of directionality, it is helpful to introduce the terms ‘detailed balance’ and ‘steady state’. For systems with at least two states, d*etailed balance* can be defined, meaning that the populations π of all states do not change over time. This even holds true if the forward and backward rates between the states are not equal. This seemingly counterintuitive statement is best explained in terms of different probabilities *p*. When looking at an example (Fig. 1) with a sparsely populated state 0 and a high transition probability (i.e. rate constant) to state 1 on the one hand, and a highly populated state 1 with a low transition probability back on the other hand, transition probability multiplied by the state population can result in the same ‘amount’ that transitions from state 0 to 1 as vice versa in a given time. This is a necessary condition for a system in detailed balance, namely that the forward ‘amount’ equals the backwards ‘amount’ (and not the rate constant) along every edge (*i* <-> *j*).

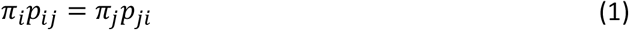

If that holds true for all edges, the populations of all states involved remain constant and are unchanged over time (stationary).

**Figure 1:**
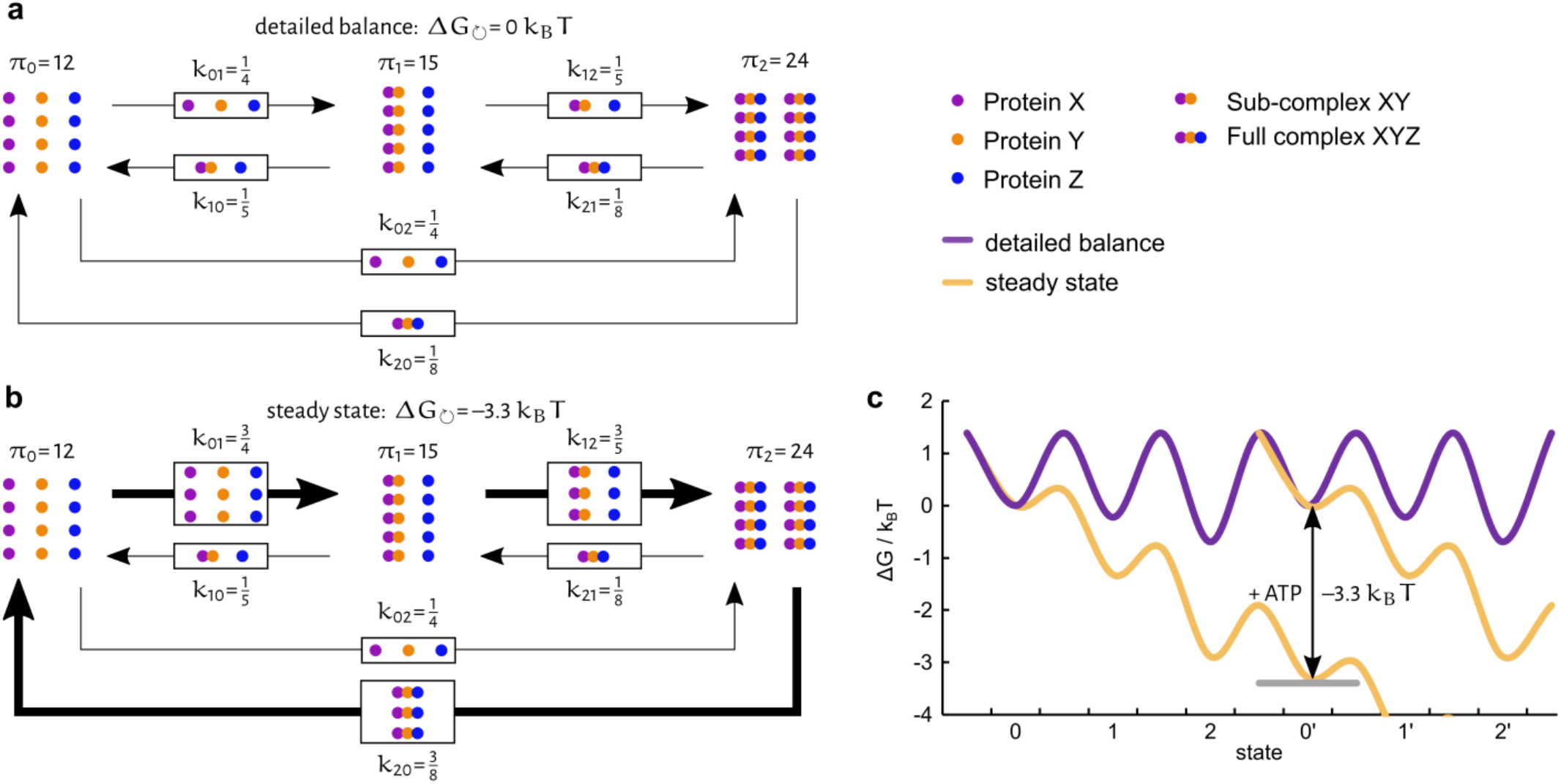
Detailed balance and steady state. Schematic representation of an exemplary system consisting of the three proteins X (violet), Y (orange) and Z (blue) in detailed balance (**a**) or steady state (**b**). Grouped protein dots represent (sub-)complexes, π_i_ signifies the state population of state *i*, the number of proteins equals the population size, and *k*_*ij*_ denotes the kinetic rates between states *i* and *j*. In detailed balance, the same amount of proteins (here: 3) change their state at each node in every time interval, although the backward and forward rates are not equal (because the state populations are not equal). In steady state, there is a net flux in one direction (here: 9 forward, 3 backward, i.e. net flux of 6), which is represented by arrow width, but the state populations stay constant, as they do in detailed balance. To obtain this directionality, energy is needed which leads to entropy production. This is depicted in the energy landscape (**c**). Here, the purple line signifies the system from (**a**) in detailed balance, while the yellow line represents the steady state condition from (**b**). In the latter case, energy (e.g. in the form of ATP hydrolysis) is needed for the system to return to the initial state and start another directed cycle. In our examples this energy is 3.3 *k*_*B*_*T*.

If there are more than two states, another possibility for stationary (constant) state populations is possible: *steady state*. In this case, e.g. more transitions occur from state 0 to state 1 than vice versa in a certain time. Those unequal forwards and backwards amounts are countered by transitions to other states of the cyclic system, allowing the population sizes of the states to stay the same. In our example (Fig. 1), we observe the binding and unbinding of three proteins X, Y, and Z. Initially, all proteins are unbound (state 0). When proteins X and Y bind, their subassembly corresponds to state 1. The resulting final complex XYZ of all three proteins corresponds to state 2. It eventually falls apart again, causing the system to return to state 0. If the system was in detailed balance (Fig. 1a), equal amounts of complexes would form and disassemble in each step. The system would be in thermodynamic equilibrium. However, with an energy source like ATP present, a succession of states could occur. In this scenario (Fig. 1b), the complex formation would be directed and a net flux through the cycle could be observed, with all forward amounts being larger than their backwards amounts. The state populations would still remain stationary (constant) but the system would be in steady state. In a cellular environment, all states are populated, and a directed cycle allows for the recycling of proteins. Steady state and detailed balance can only be clearly distinguished with single-molecule experiments, because from an ensemble view, all state populations stay constant in both cases.

Another possibility to introduce directionality would be the successive binding of proteins by utilizing the energy of binding. However, this can never lead to a cycle as for every forward transition, a new protein would have to be synthesized to be pristine for binding. In the end, the very stable complex would have to be degraded because it constitutes the energy minimum of the binding process. This single-use of proteins would result in a huge energy cost and is therefore very unlikely.

Mathematically, directionality can be expressed in terms of Gibb’s free energy ΔG_↺_, i.e. the entropy production during one cycle, being different from 0.

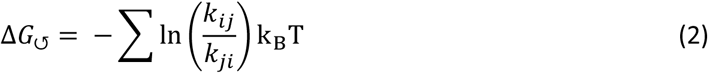

In this equation, *k*_*ij*_ is the forward rate from state *i* to state *j, k*_*ji*_ is the corresponding backward rate, k_B_ is the Boltzmann constant and T is the temperature.

As mentioned above, directional cycles have often been suggested for the Hsp90 machinery, linking its conformational states to its ATPase function. Experimentally, several conformational states have been observed by x-ray crystallography and cryo-EM (Verba et al. 2016; Ali et al. 2006; Bron et al. 2008). The dynamics between these conformational states can be measured with single-molecule Förster Resonance Energy Transfer (smFRET), similar to many other protein systems (Tassis et al. 2021; Evans et al. 2022; Krainer et al. 2020; Steffen et al. 2020). From the FRET efficiency, an open and closed conformation can be distinguished. Although its opening and closing dynamics suggest a two-state system, Hsp90 actually occupies at least four states, displaying two degenerate (with respect to FRET efficiency) open states (0 and 1) and two degenerate closed states (2 and 3) which differ kinetically. A separation of the degenerate pairs, however, requires the analysis of their kinetics. Previously, we have shown that these four states are hardly coupled to ATP hydrolysis for the idle Hsp90, i.e. in the absence of any other proteins (Mickler et al. 2009; Schmid et al. 2016).

Even though ATPase activity seems to be a prerequisite for client maturation, yeast Hsp90 displays a slow ATPase rate, only hydrolyzing about one ATP per minute (Panaretou et al. 1998). In this publication, we added several co-chaperones (Aha1, Sba1, Cdc37) and the client kinase Ste11 to understand when the energy of ATP hydrolysis is coupled to Hsp90’s conformational changes and therefore, when directionality is introduced. All of the cochaperones used within this paper are known to modulate Hsp90’s ATPase activity: while Aha1 enhances it (Wolmarans et al. 2016; Panaretou et al. 2002), Sba1 (McLaughlin et al. 2006; Richter et al. 2004; Siligardi et al. 2004) and Cdc37 have been shown to inhibit it (Eckl et al. 2013). In this context, Sba1 has been found to stabilize client-activating states, preferably binding to ATP-bound Hsp90 (Ali et al. 2006; Ratzke et al. 2014). Cdc37, on the other hand, has been dubbed a ‘kinase-linking cochaperone’ due to its specific and dynamic interactions with inactive kinases and Hsp90 (Keramisanou et al. 2022; Shao et al. 2001; Eckl et al. 2015; Verba et al. 2016). Here we confirm that the presence of Cdc37, the kinase Ste11 and ATP lead to changes of Hsp90’s kinetic rates, but surprisingly they do not introduce directionality. However, once Aha1 and Sba1 are added, clear directionality can be observed. Therefore, all three cochaperones are necessary to convey the energy of ATP hydrolysis.

## Results

### Directionality can be detected and quantified with single-molecule FRET experiments

To observe and quantify a system’s directional behavior on a single-molecule level, data acquisition, analysis and thereupon modelling require high levels of reliability – especially when the expected changes in Gibb’s free energy are small. In fact, this has only been done for proteins that move either linearly like kinesin or myosin (Svoboda et al. 1993; Sheetz et al. 1984), or rotate like the F_0_F_1_-ATPase (Yasuda et al. 2001).

To address the challenge of observing directionality in conformational changes, we have developed a new testing procedure that uses external laser triggering to create artificial directionality in experimental data. This method was tested using a Total Internal Reflection Fluorescence (TIRF) microscope for data acquisition and Hidden Markov Modeling for analysis and evaluation. The artificial directionality was introduced by laser triggering, creating a loop of four different states with a defined sequence (Fig. 2a): a long-lived high-FRET state followed by a long-lived low-FRET state, then a short-lived high-FRET state before a short-lived low-FRET state. While the lifetime of the states was controlled by illumination time, the difference in FRET efficiency was achieved by using an additional laser. When labelled low FRET dsDNA (Hellenkamp et al. 2018) is simultaneously excited by the donor (green) and the acceptor (red) laser, it artificially displays “high FRET” by direct excitation. In contrast, the dsDNA shows “low FRET” if only excited with the green laser.

**Figure 2:**
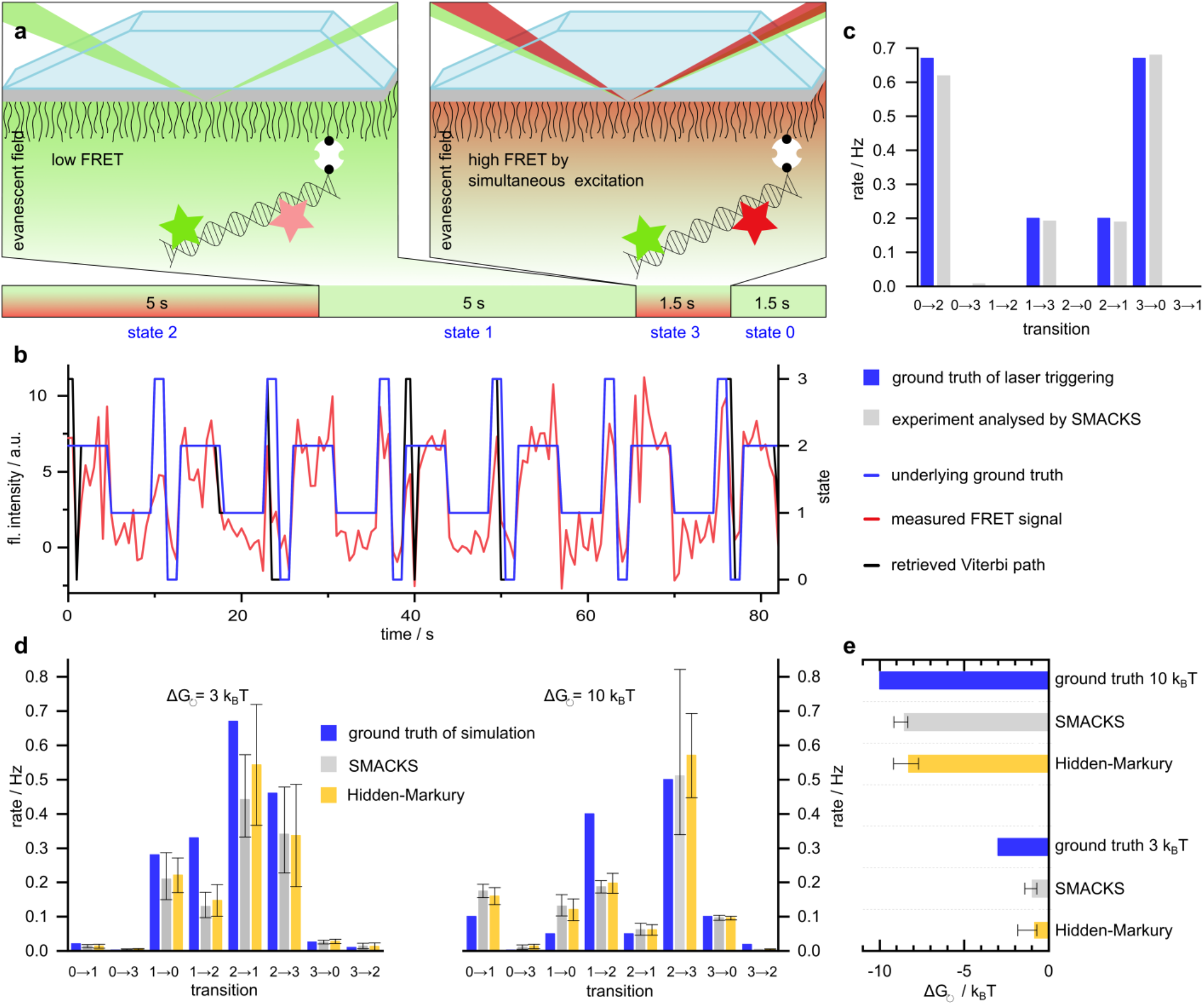
Our single-molecule detection and analysis is capable of quantifying directionality. (**a**) Artificially introduced directionality by laser triggering. To achieve two distinct FRET efficiencies, a fixed low-FRET dsDNA sample is illuminated alternatingly with a green laser only (low FRET) or with a red and a green laser simultaneously (artificial high FRET). Lifetime of the states can be controlled through different illumination times (1.5 vs 5 seconds). (**b**) Pattern of the laser triggering as programmed (underlying ground truth, blue) as well as the recorded experimental data (FRET signal, red trace). The Viterbi path was retrieved by data analysis with SMACKS and is shown in black. The results of the kinetic rates are shown in (**c**). The experiment is in very good agreement with the laser induced ground truth. (**d**) Ground truth of data simulation for two different values of directionality (blue, 3 k_B_T vs 10 k_B_T) and the results from analysis with SMACKS (grey) and the Hidden-Markury software (yellow). The transition rates are mostly in good agreement. (**e**) Retrieved Gibb’s free energy from the analysis of the simulated data (ground truth, blue) with SMACKS (grey) and Hidden Markury (yellow). The higher the given net flux, the better the Gibb’s free energy can be retrieved by HMM based software.

Fig. 2b and 2c show that the ground truth and experimental model are in very good agreement. The deviations of the populations are less than 4 %, while the transition rates deviate by a maximum of 8 %. Although only a forward progression through the cycle was triggered, the analysis retrieved a few false backwards rates due to wrong state allocation. The resulting very small backward rates allowed for a ΔG calculation amounting to - 165 k_B_T. In contrast, the calculation of a ΔG for the ground truth is mathematically not defined as there are no backwards rates. Altogether, these results prove that directionality can be detected and analysed on the single-molecule level.

To quantify the directionality (entropy production) that can be retrieved by our software SMACKS (Schmid et al. 2016) and to compare it to another software (Hidden Markury (Gebhardt 2021)), we used simulated data with a directionality of 3 k_B_T and 10 k_B_T, respectively. The data was simulated with MASH FRET (Börner et al. 2018) and then analysed with SMACKS and Hidden Markury. Both programmes, which have previously been tested in a comparative study (Götz et al. 2022), were able to retrieve the pre-specified ΔGs, although errors became bigger for smaller ΔG (Fig. 2D and 2E). For ΔG=3 k_B_T, both software programmes resulted in about 30 % of the ground truth, whereas for ΔG=10 k_B_T about 85 % were recovered. In both cases, the directionality was underestimated, which is expected, as the entropy production from Hidden Markov Models (HMM) is only a lower limit (Godec and Makarov 2023).

### Directionality in the Hsp90-kinase machinery requires three cochaperones

After having ensured the robustness of both data acquisition and analysis, our experiments were extended to the Hsp90 machinery. Fig. 3a,c shows examples of dynamic smFRET traces of Hsp90’s opening and closing dynamics in presence and absence of cochaperones and a client kinase. The FRET pair positions chosen on Hsp90 (61 and 385, one on each monomer) are well-established for these experiments, since they show a substantial difference between Hsp90’s two closed high FRET states and its two open low FRET states (Schmid et al. 2016; Schmid and Hugel 2020; Mickler et al. 2009). Both high FRET as well as both low FRET states are degenerate in their FRET efficiency but can be separated by their kinetics (Schmid et al. 2016).

**Figure 3:**
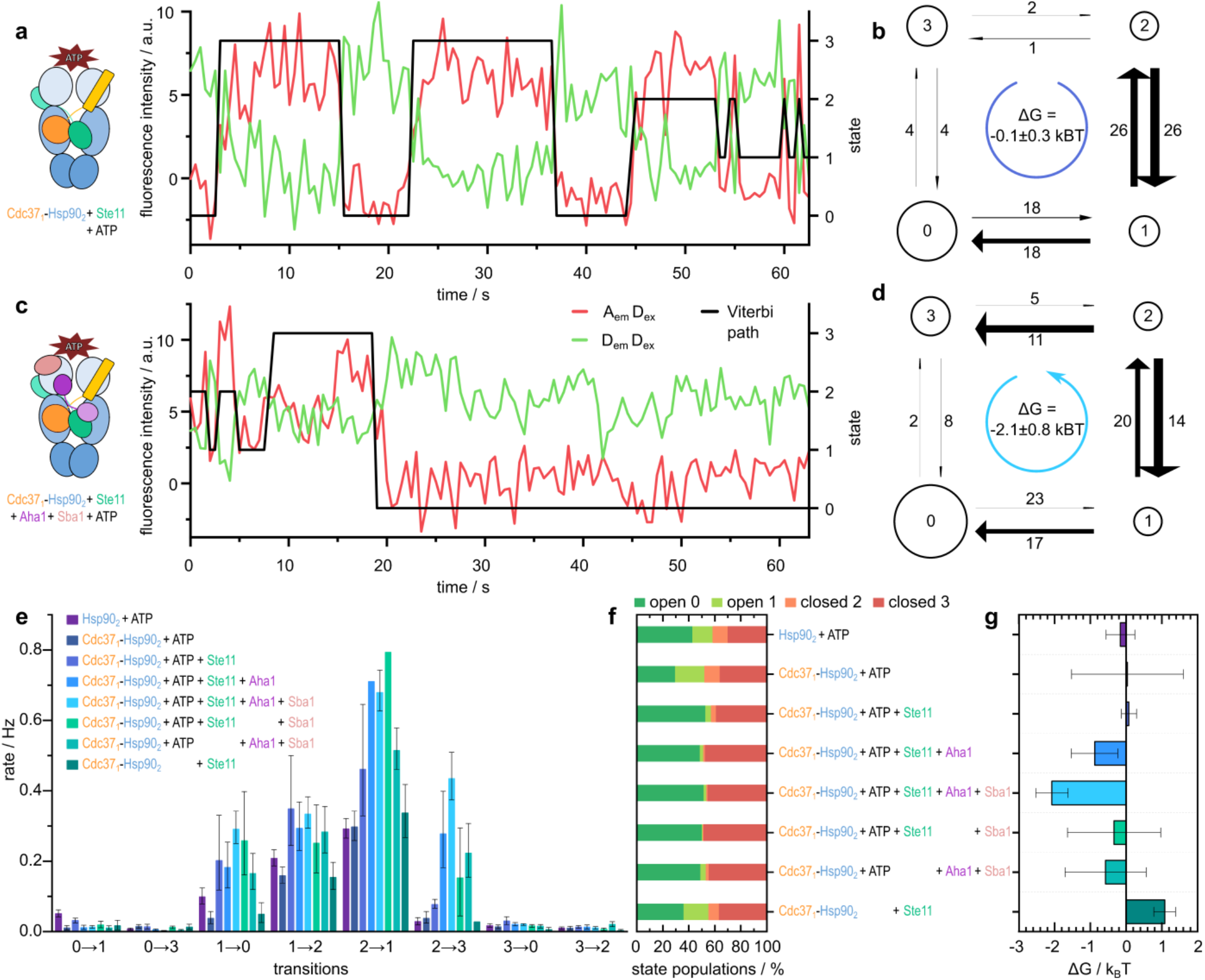
Three cochaperones are needed to induce directionality. Hidden Markov analysis of single-molecule traces (**a, c**) determines the Viterbi paths (black line), the transition rates (**e**) and the state populations (**f**), which can be represented as models (**b, d**) and allow for calculation of the entropy production ΔG (**g**). **(a, b**) Cdc37_1_-Hsp90_2_ + ATP + Ste11 in detailed balance: a directed progression through Hsp90’s conformational states can neither be observed in the Viterbi path (**a**) nor the corresponding model (**b**) (n = 509 traces). (**c, d**) Cdc37_1_-Hsp90_2_ + ATP + Ste11 + Aha1 + Sba1 in steady state: single-molecule trace displays a directed Viterbi path through all states, starting with fluctuations between state 1 and 2, then transitioning to long closed (state 3) and afterwards to long open (state 0) (n = 998 traces). (**b, d**) Four-state models with equal FRET efficiencies for states 0/1 (open) and 2/3 (closed), respectively. Circle sizes are proportional to the state population, arrow widths to the transition rates. Numbers on arrows indicate normalized amount of transitions occurring. (**e**) Overview of transition rates for each measured condition. The presence of all three cochaperones (lightest blue) leads to the strongest increase in rate 2→3. (**f**) Percentage of state population for each measured condition. The presence of client and cochaperones causes longer dwell times in the long-lived states 0 and 3. (**g**) Overview of calculated Gibb’s free energy for each measured condition. Entropy production can only be observed in presence of all three cochaperones (lightest blue), the condition without ATP (dark green, bottom) was used to determine an error for ΔG. Results shown in (**e**-**g**) are obtained from two to four independent repetitions with 354 to 998 traces in total.

Here we refer to Hsp90’s open states as state 0 and 1, and its closed states as state 2 and 3. In general, forward and backward rates for transitions between all of them can be obtained by kinetic analysis with SMACKS (Schmid et al. 2016). However, diagonal rates (02, 20, 13 and 31) were always close to zero and are therefore not considered in the following.

First, we tested if Cdc37, Ste11 and nucleotide introduce directionality, which is the currently prevailing model (conditions with one Cdc37 per Hsp90 dimer (Cdc37_1_-Hsp90_2_) were used). Fig. 3a shows an exemplary smFRET trace and Fig. 3b the extracted model. Note that the Viterbi path (black trace) is not a fit to the data, but the model extracted from the data, with the ‘state’ axis on the right. Fig. 3e shows clear changes compared to idle Hsp90 in the transition rates 10, 01 and 12. Rate 21 is increased as well, but within the 95 % confidence intervals. However, despite noticeable rate changes, the system stays in detailed balance, i.e. there is no directionality (Fig. 3e and 3g).

Therefore, we tested if the addition of further cochaperones – Aha1 and Sba1 – to the system introduce a directed progression of Hsp90 through its conformational states (Fig 3c and 3d). Indeed, in this case not only the transition rates are changed, but the system also shows a Gibb’s free energy of Δ*G* ≈ -2.1 k_B_T, meaning that entropy is produced and the system is no longer in detailed balance. This behavior is mostly due to a strong increase in rate 23 (Fig. 3e and 3g).

As the such calculated free energy is a lower bound for the energy coupled into the open-close movement, this is a first evidence that the energy of ATP hydrolysis couples to the large conformational changes of Hsp90, although only in presence of three cochaperones and a substrate. In further experiments, the effect of either Aha1 or Sba1 on the minimal model was tested as well, however, no directionality was induced by the sole addition of one interaction partner. This indicates that the Hsp90 machinery relies on a larger number of interaction partners to escape detailed balance and enter steady state and therefore to convey the energy of ATP hydrolysis.

In contrast to the detectable changes in its kinetic rates, a look at Hsp90’s state populations reveals that none of the examined conditions significantly affected thermodynamics: Hsp90‘s different interactors do not initialise a closing of the dimer, i.e. the ratio of open to closed Hsp90 was not strongly altered (Fig. 3f). This is especially striking for the presence of Cdc37, as it inhibits Hsp90’s ATPase rate (Fig. S1). This again indicates that Hsp90’s conformational state is hardly connected to ATP hydrolysis. Even more astounding: Aha1’s induced closing of the dimer (Schulze et al. 2016) is abolished in presence of Cdc37 and Ste11.

The presence of Cdc37, Ste11 and ATP (both in presence and absence of Sba1 and Aha1), ‘locks’ almost half the complexes in the longer-lived states 0 and 3, respectively. This can be explained by the strongly increased rates 10 and 23. The enhancement of the opening rate 21 adds to this as well but is compensated for by the simultaneous rise of its opposed closing rate 12, meaning that the exchange between the short-lived states is accelerated.

Altogether, the changes in Hsp90’s kinetic rates indicate a cooperative interplay between Cdc37 and Ste11 in presence of ATP. These changes, however, are not yet sufficient to introduce directionality. For an ATP-driven cycle of Hsp90’s conformational kinetics – and therefore cyclic operation of Hsp90 – additional Aha1 and Sba1 are required, suggesting their active role in conveying the energy provided by ATP. Please note that the alternative to a cyclic operation is an energetic downhill process through successive binding of proteins. This is very unlikely because it would mean that one Hsp90 has to be degraded at the end of the energetic downhill process for each kinase activation event.

### Directionality and ATPase rates are only weakly linked in Hsp90-kinase complexes

In the following, we tested how the directed progression of Hsp90’s dynamic, i.e. cycling, is linked to Hsp90’s ATPase rate. Therefore, we determined Hsp90’s ATPase in presence of Cdc37, Sba1, Aha1, Ste11 and their combinations. Hsp90’s ATPase rate was found to be 1.3 ± 0.3 ATP/minute, fitting well to previous data (Panaretou et al. 1998). Fig. 4 shows that in presence of equimolar amounts of only Cdc37 and Ste11 – a condition that did not induce directed behavior of Hsp90’s kinetics – the chaperone’s ATPase rate is not significantly altered. This implies that there is an interaction between the kinase and Cdc37, preventing the latter from performing its full inhibitory function (Fig. S1). However, the presence of Aha1 enhances this rate in all of the following cases, albeit to a different degree: when Cdc37 is present as well, the rate is increased to 1.8 ± 0.2 ATP/minute. When Ste11 is added to the mix, it is further enhanced to 2.2 ± 0.2 ATP/minute. Surprisingly, upon the addition of Sba1 – a known ATPase inhibitor – Hsp90’s ATPase rate is boosted further to 2.6 ± 0.3 ATP/minute. This is in good agreement with the directionality observed in single-molecule measurements, indicating that there is some interplay that connects the components’ interaction to Hsp90’s ATP hydrolysis. Altogether, such an interaction between three cochaperones allows for more evolved regulation as e.g. through a typical nucleotide exchange factor.

**Figure 4:**
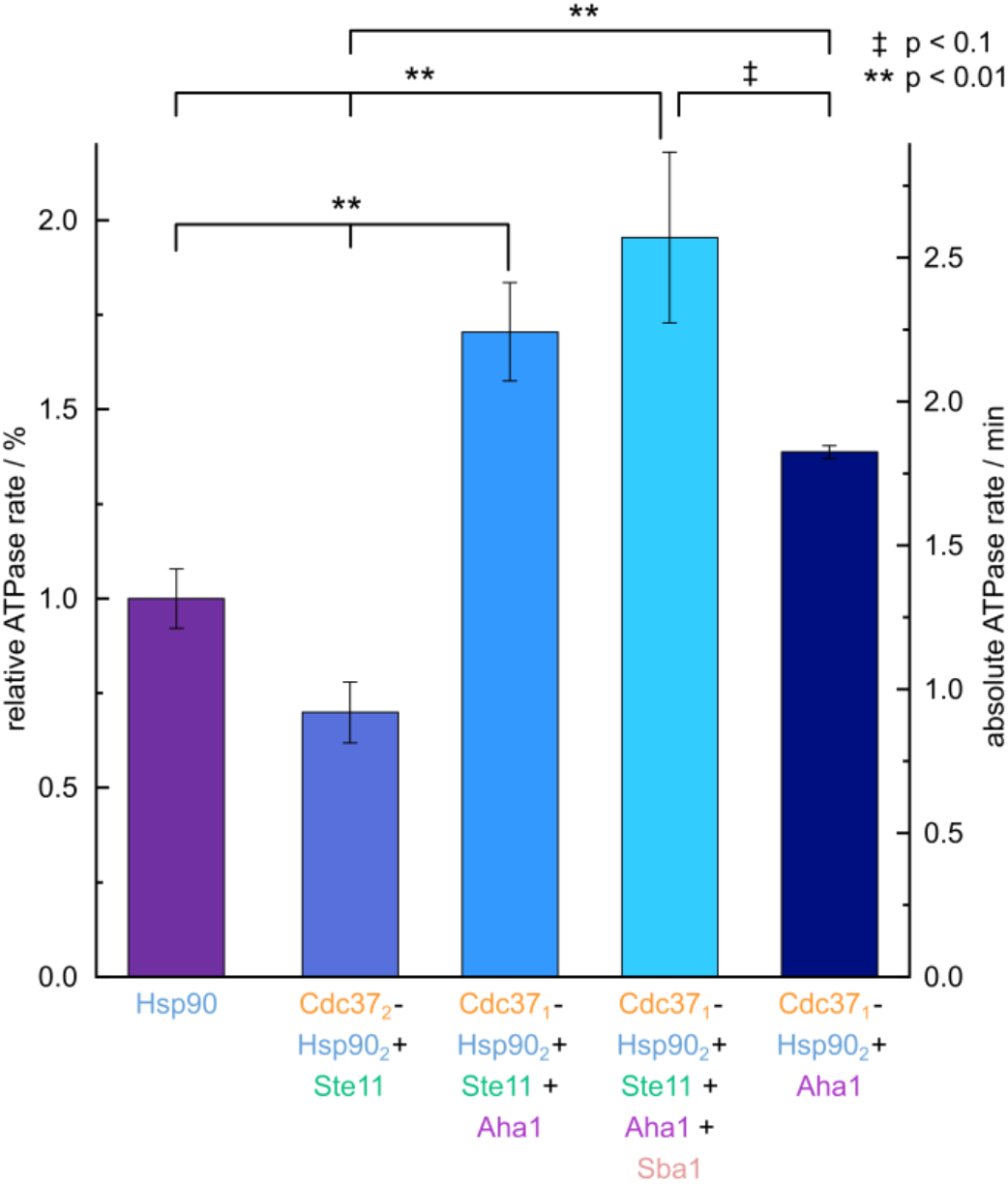
Relative and absolute ATPase rates of Hsp90 in presence and absence of the cochaperones Cdc37, Aha1 and Sba1 and the client kinase Ste11. Hsp90 is a slow ATPase with 1.3 hydrolysed ATP per minute (violet). Ste11 hinders Cdc37’s ATPase-decreasing effect on Hsp90 (blue, second to left bar). Aha1, Cdc37, Sba1 and Ste11 together (light blue, second to right bar) strongly increase the ATPase to 2.6 ATP/min. This effect is not achieved by Cdc37 and Aha1 alone (darkest blue). Statistical significance tested by one-way ANOVA and Tukey post hoc test (described in methods). Additional conditions are shown in Fig. S1.

## Discussion

Most suggested conformational cycles of Hsp90 imply strong directionality as they are commonly depicted as being unidirectional. Here we show a fundamentally different scenario, in which an upstream equilibrium allows for an assembly of most of the involved proteins, and then one (or few) directed steps stabilize the functional complex (Fig. 5). We consider this scenario to be much more likely for the formation of any multi-protein complex for several reasons. First, for an energetic reason: Consider a complex of five molecules with a defined binding order. To achieve a reasonably defined binding sequence, the energy of at least one ATP hydrolysis (around 20 k_B_T) needs to be consumed in each step. This could for example be provided as binding free energy. However, this would eventually result in a very stable complex that could only be broken apart again upon an energy input corresponding to five ATPs – one for every step – or by degradation of some of the proteins involved. In our opinion, both options seem unlikely, as they would be highly inefficient and a Hsp90 dimer binds (and hydrolyses) two ATP at maximum per cycle. In contrast, an upstream equilibrium as suggested here would only involve energies of a few k_B_Ts. Final formation of the functional complex would then involve only stabilizing energy of e.g. one ATP hydrolysis. Therefore, the components could easily be recycled by the hydrolysis of one or two ATP, which are available from Hsp90.

**Figure 5:**
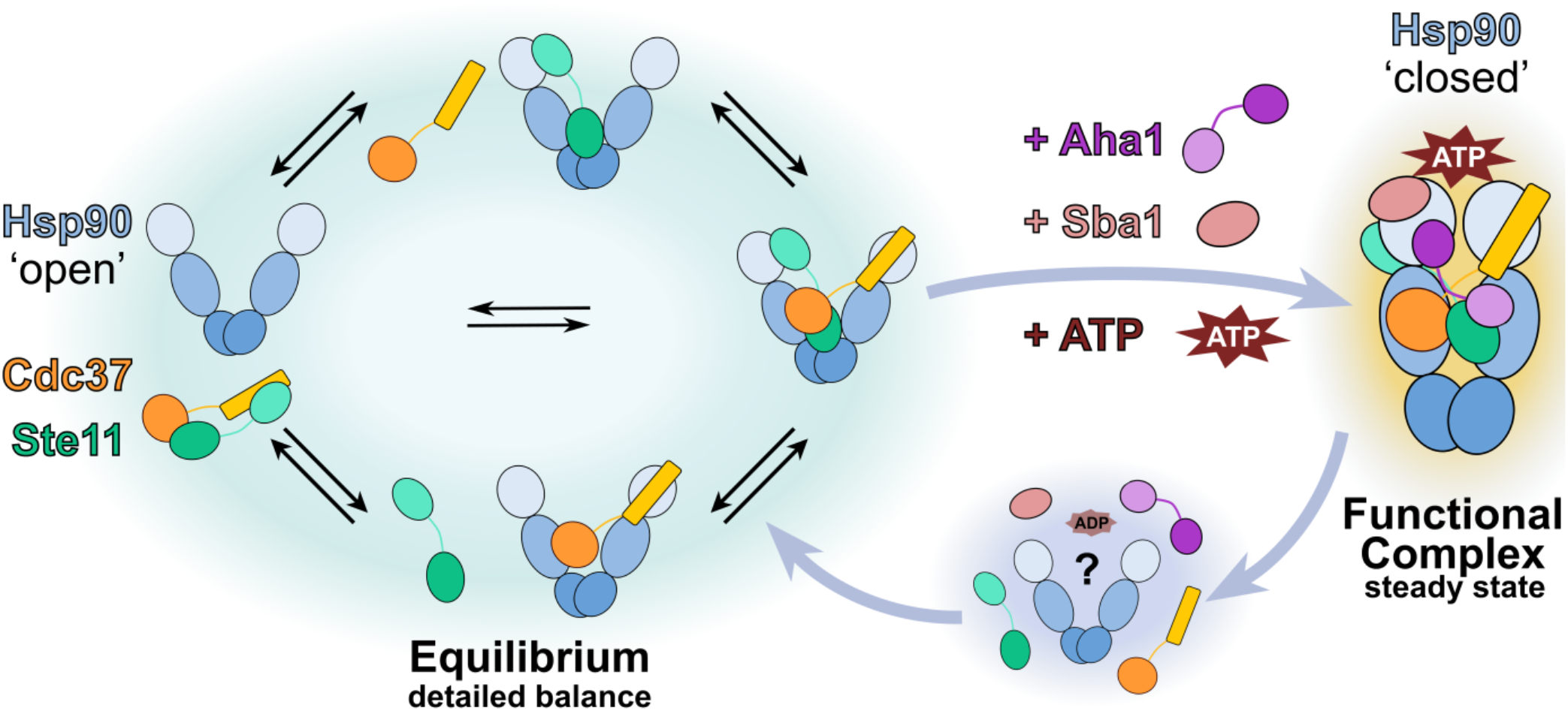
From sub-complex assemblies in equilibrium (green halo) to the functional multi-protein-Hsp90 complex (orange halo). Sub-complexes of Hsp90, Cdc37 and Ste11 are in detailed balance, thus assemble and disassemble in equilibrium. Upon the addition of Aha1, Sba1 and ATP, the functional complex is stabilized, e.g. by utilizing the energy provided by ATP hydrolysis. Therefore, recycling of the components and complex disassembly (blue halo) would only require the energy provided by hydrolysis of only one or two ATPs. The third state where all proteins are unbound (blue halo) was not quantified and could also be part of the left equilibrium, but more than two states are necessary for directionality (blue arrows). Another possibility for this third state could be a partially disassembled state with hydrolysed ATP (ADP).

A second reason is the timescale of complex formation: If this were a strongly directed downhill process, it could easily happen in a fraction of a second (motors like myosin, kinesin or the F_0_F_1_-ATPase hydrolyse ATP with up to several hundred per second). This, however, is not necessary in our opinion. Dynamic complex formation on the timescale of seconds allows for much more regulation, e.g. by post-translational modifications (PTMs). Third, our data clearly shows that there is no linear succession of binding events. To our knowledge, there is also no published study, that shows such a succession. Of course, if components are added in a certain order, an artificial succession is introduced, however, this does not serve as evidence for succession of binding in steady state or even living cell conditions. Actually, such an artificial introduction of proteins would correspond to protein production and degradation in every cycle. Up to this point, we even have neglected conformational changes, which define multi-conformational complexes (Hiller 2019). It is important to note that not only Hsp90 samples different conformations, but also its cochaperones. Their thorough study is therefore necessary to complete understanding of the machinery. Cdc37, for example, was shown to change between a dynamic extended conformation and a compact structure to sense partially unfolded clients and then stabilizing them (Keramisanou et al. 2022). In this context, a directional cycle might also be obtained by looking at yCdc37_S14’s dephosphorylation by protein phosphatase 5 (PP5 in humans, Ppt1 in yeast), which is thought to be important for kinase release (Oberoi et al. 2016; Sager et al. 2020).

If these conformational changes and PTMS were also included into the successive, directed assembly, even more energy would be required. In addition, we and others have shown before that Hsp90’s conformational changes occur on much faster timescales than ATP hydrolysis (Mickler et al. 2009; Schmid et al. 2016). Thus, conformational changes are thermally driven and not fueled by ATP turnover. A recent study suggests that it is not the hydrolysis of ATP but the repositioning of its gamma phosphate which is sufficient for Hsp90 function (Reidy et al. 2023). They emphasize that returning to the open, ADP bound state is essential, which can occur either by nucleotide exchange or by hydrolysis. ATP hydrolysis is evolutionary preferred because it allows for a regulatory switch that can be tuned by cochaperones. By regulating the ATPase activity, they regulate the dwell times of Hsp90’s conformational states. In agreement with this observation, we see a shift of dwell times to the long-lived conformational states in our single-molecule experiments (Fig. 3f).

Finally, our data is in perfect agreement with an assembly mechanism as depicted in Fig. 5. No directionality can be detected in any sub-system of the five components Hsp90, Cdc37, Aha1, Sba1, Ste11, i.e. they are close to thermal equilibrium. Only when all five components are together and ATP is present, directionality can be detected. This would allow first for the formation of a functional complex and then, e.g. after ATP hydrolysis or nucleotide exchange (Reidy et al. 2023), for recycling of all components. We speculate that this is the long searched for function of Hsp90’s ATPase function, namely to disassemble a functional complex after the job is done. Checkpoints might be included without the need of large amounts of free energy.

## Acknowledgements

This work was supported by the Deutsche Forschungsgemeinschaft (DFG) under Germany’s Excellence Strategy (CIBSS EXC-2189 Project ID 390939984) and the SFB1381 programme (Project ID 403222702). We thank Aljaz Godec and Dimitrii Makarov for helpful discussions. We thank Marianne Birkle and Michael Witt for support with protein purification.

## Author contributions

T.H. designed the research; J.S., L.V. performed the measurements; B.H. made the protein constructs. J.S., L.V. analysed the data after consultation with T.H. All authors wrote the manuscript and have given approval to the final version of the manuscript.

## Methods

### Expression constructs

All proteins used in this study originated from *Saccharomyces cerevisiae*.

Hsp90 constructs contained the *hsp82* gene from *Saccharomyces cerevisiae*, an N-terminal cleavable His_6_-SUMO-tag and a C-terminal coiled-coil domain ensuring stable dimer formation (Mickler et al. 2009). The coiled-coiled motif is either followed by a Strep-tagII or an Avi-tag for *in vivo* biotinylation.

The basic Cdc37-Hsp90 fusion protein was purchased as synthetic gene construct (GeneScript). Such fusions provide for a high local concentration and overcoming low binding affinities in single moelcule experiments (Siligardi et al. 2002). The linker between the two proteins consists of in total 115 amino acids, with the following sequence: P-(GGS)_15_-P(EAAAK)_3_-P-(GGS)_15_-P (Chen et al. 2013). The construct also carried the C-terminal coiled-coil motif followed by either a Strep-tagII or Avi-His_6_-tag for *in vivo* biotinylation.

The Cdc37 domain from the fusion-protein additionally carried two point mutations, S14E and S17E, the first glutamate mimicking a phosphorylated serine residue, the second was introduced due to the presence of a natural glutamate forming a stabilizing salt bridge, as derived from a Cryo-EM complex structure (Verba et al. 2016). In the free Cdc37 only the S14E phosphomimic was present.

All expression constructs were cloned into pET28b plasmid vectors and single cysteines in Hsp90 at position 61 or 385 were introduced in every construct for site-specific labelling. Free Aha1, Sba1 and Cdc37 constructs carried a N-terminal His_6_-tag for purification.

### Protein preparation

Expression and purification was performed in variations of established protocols as previously reported (Tych et al. 2018; Schmid and Hugel 2020). In short, proteins were produced in *E*.*coli* BL21Star (DE3) or BL21 (DE3) cod+ and purified depending on the affinity tag. For in *in vivo* biotinylation on the Avi-tag the plasmid pBirA (Avidity Nanomedicines, La Jolla, CA) was cotransformed and the cell culture procedure was adapted according to Avidity’s *in vivo* biotinylation protocol.

His_6_-SUMO tagged Hsp90 was isolated with affinity chromatography using a Ni-IMAC column (HisTrap HP), followed by SUMO cleavage with SenP protease and a second Ni-IMAC to separate the His-tag free Protein from uncleaved proteins and the His_6_-SUMO-tag. The protein was then applied to an anion exchange column (HiTrap Q, Cytiva) and finally to a size exclusion column (Superdex 200, Cytiva).

Strep-tagII and simple His_6_-tag preparations were performed with a two-step protocol, starting with affinity chromatography using either a Strep-Tactin column (Strep-Tactin Superflow, IBA Lifesciences) or an Ni-IMAC (Cytiva/ Ni-NTA Agarose, Qiagen) followed by Size exclusion chromatography (Superdex 200, Cytiva).

### Protein Labelling and monomer exchange

Hsp90_2_ and Cdc37_1_-Hsp90_2_-fusion were labelled with maleimide fluorescent dyes (ATTOTEC), as they allow for specific attachment to the introduced cysteines. First, the respective protein sample was incubated with 10 mM tris(2-carboxyethyl)phosphine (TCEP) in a volume of 50 μl and a concentration of 100 μM protein. After 20 minutes of incubation at room temperature, TCEP was removed by washing the sample five times with PBS buffer (100 mM, pH 6.7) in a centrifugal filter (Amicon Ultra, cutoff 30K). The sample was then incubated, using either Atto550 or Atto647N (1.5 times molar excess). After two hours at room temperature, free dye was removed using a spin column (PD 25 MiniTrap, GE Healthcare) preequilibrated with measurement buffer (40 mM HEPES, 150 mM KCl, 10 mM MgCl_2_, pH=7.4). The labelling efficiency was determined using a NanoDrop (Thermo Scientific) and usually reached between 80 and 100%. As the dynamic rates between two point mutations of Hsp90 (D61C and Q385C) were monitored, monomer exchanges were performed by incubating a mixture of both variants for 40 minutes at 43 °C and 300 rpm in a shaking incubator (Thermomixer 5436, Eppendorf). This method was also employed to obtain Cdc37_1_-Hsp90_2_-fusion heterodimers (one Cdc37 per Hsp90 dimer). In case of the single-molecule experiments, the mixtures contained a ratio of 1:2 of biotinylated to non-biotinylated protein, as for these measurements only the biotinylated heterodimers would bind to the chamber surface. That way, the biotinylated protein has a higher chance of being exchanged with a non-biotinylated variant. For the ATPase assays, the exchange ratio between both variants was 1:1 resulting in a binomial distribution of one part Cdc37_2_-Hsp90_2_, two parts Cdc37_2_-Hsp90_2_, and one part Hsp90_2_.

### Single-molecule measurements

The measurements were conducted at a custom-built prism type TIRF setup including two lasers of wavelength 532 nm (green, Coherent OBIS LS) and 637 nm (red, Coherent OBIS LX) as introduced in (Götz et al. 2018). The laser beam is focused perpendicularly onto a prism, which is located on top of the measurement chamber. To avoid unspecific protein binding to the flow chamber, its surface is passivated with an 80:3 mixture of methoxy-polyethylene glycol-silane (5000 Dalton, Rapp Polymere) and silane-PEG-Biotin (3000 Dalton, Rapp Polymere). To further enhance this passivation, the measurement is preceded by 30 minutes of BSA incubation (0.5 mg/mL in measurement buffer, Carl Roth). The biotinylated protein sample is attached to the surface via biotin-Neutravidin binding. Therefore, the chamber was further coated with Neutravidin (0.25 mg/mL, Thermo Fisher Scientific) previous to the addition of the sample (picomolar range). Free cochaperones (Aha1 and Sba1) and Ste11 (*Saccharomyces cervisiae* Ste11-(AA 415-712)-P23561-partial protein, Cusabio, washed with measurement buffer) were added at 2 μM each with 2 mM ATP present.

The sample was measured in alternating laser excitation (ALEX) mode between the green and the red laser, with excitation times of 200 ms and dark times of 50 ms. Fluorescence signal was collected by an oil immersion objective (100x magnification, Nikon) and recorded by EMCCD cameras (iXon Ultra 897, Andor) at 3x3 binning. Image registration with fluorescent beads (TetraSpeck microspheres, 0.2 μm, Invitrogen) preceded the experiments to align the green and the red channel.

An Igor Pro (version 6.37) based in-house script was used for the selection of single-molecule traces. The program identifies the positions of single-molecules by searching for the brightest spots in five consecutive frames of the respective detection channel.

### Laser triggering for artificial directionality

Artificial directionality for two-colour single-molecule Förster Resonance Energy Transfer (FRET) was induced by laser triggering and recorded in ALEX mode. The repeating loop consisting of a short-lived high-FRET state followed by a short-lived low-FRET state, then a long-lived high-FRET state before a long-lived low-FRET state was programmed in LabView 2019. The length of both short-lived states was set to 3 consecutive frames, whereas for both long-lived states, a duration of 10 frames was chosen. The difference in FRET efficiency was achieved by using a labelled dsDNA sample displaying a low FRET efficiency (E = 0.15 ± 0.02, 1-lo sample (Hellenkamp et al. 2018), biomers.net GmbH Ulm) and switching between excitation by the green laser only (low FRET) and simultaneous excitation by the green and the red laser (high FRET).

### Simulation of dynamic time traces

Simulation of dynamic time traces was done with MASH-FRET (Börner et al. 2018). For all simulations, four FRET states with two degenerate low as well as two degenerate high FRET efficiencies (E_low_= 0.1 ± 0.05, E_high_= 0.8 ± 0.05) were given. The transition matrix for a Gibb’s free energy of 3 k_B_T and of 10 k_B_T were predefined (Tab. 1). 200 traces with 200 frames with a frame rate of 2 Hz were simulated, respectively.

**Table 1.** Parameters for trace simulation

| Ground truth with $\Delta G = 3 k_B T$ | | | | |
| --- | --- | --- | --- | --- |
| FRET E | $E_0 = 0.1$ | $E_1 = 0.1$ | $E_2 = 0.8$ | $E_3 = 0.8$ |
| kinetic rates [Hz] | | $k_{01} = 0.02$ | $k_{02} = 0$ | $k_{03} = 0.002$ |
| | $k_{10} = 0.28$ | | $k_{12} = 0.33$ | $k_{13} = 0$ |
| | $k_{20} = 0$ | $k_{21} = 0.67$ | | $k_{23} = 0.46$ |
| | $k_{30} = 0.025$ | $k_{31} = 0$ | $k_{32} = 0.01$ | |
| Ground truth with $\Delta G = 10 k_B T$ | | | | |
| FRET E | $E_0 = 0.1$ | $E_1 = 0.1$ | $E_2 = 0.8$ | $E_3 = 0.8$ |
| kinetic rates [Hz] | | $k_{01} = 0.1$ | $k_{02} = 0$ | $k_{03} = 0.002$ |
| | $k_{10} = 0.05$ | | $k_{12} = 0.4$ | $k_{13} = 0$ |
| | $k_{20} = 0$ | $k_{21} = 0.05$ | | $k_{23} = 0.5$ |
| | $k_{30} = 0.1$ | $k_{31} = 0$ | $k_{32} = 0.018$ | |

### Single-molecule data analysis

The dynamic analysis of both simulated and experimentally measured data was done using two programmes employing Hidden Markov Models. In SMACKS (Schmid et al. 2016), an Igor Pro (version 6.37) based script, two apparent states were chosen for FRET state assignment. A four-state model with two hidden states was chosen by Bayesian information criterion (BIC) and subsequently a transition matrix was calculated. In Hidden Markury, a python-based notebook (Gebhardt 2021), the same four-state model was assumed and diagonal rates were set to zero. For rate calculation, the 2D model was used, which retrieves the input from donor and acceptor signal. Mean transition rates were calculated for each condition from *n* replicates (Tab. 2). 95% confidence intervals (CI) resulting from each HMM modelling were treated as standard error of the means and averaged as such:

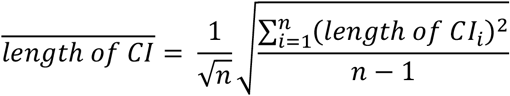

To obtain an independent estimate for the error of the determined Gibb’s free energy, control experiments involving Hsp90 in presence of Cdc37 and Ste11 without ATP were measured. This minimal model lacking an energy source displayed a ΔG of + 1.1 k_B_T.

**Table 2.** Number of repetitions and amounts of analysed single-molecule traces.

| Condition | Number of replicates | Total number of traces |
| --- | --- | --- |
| Hsp90 + ATP | 4 | 896 |
| Cdc37<sub>1</sub>-Hsp90 + ATP | 3 | 640 |
| Cdc37<sub>1</sub>-Hsp90 + ATP + Ste11 | 2 | 509 |
| Cdc37<sub>1</sub>-Hsp90 + ATP + Ste11 + Aha1 | 2 | 614 |
| Cdc37<sub>1</sub>-Hsp90 + ATP + Ste11 + Aha1 + Sba1 | 3 | 998 |
| Cdc37<sub>1</sub>-Hsp90 + ATP + Ste11 + Sba1 | 2 | 449 |
| Cdc37<sub>1</sub>-Hsp90 + ATP + Aha1 + Sba1 | 2 | 527 |
| Cdc37<sub>1</sub>-Hsp90 + Ste11 | 2 | 354 |

### ATPase assays

The ATPase activities of the protein samples were measured with a UV/Vis Spectrometer (Lambda 35, Perkin Elmer) using an ATP (2 mM, Jena Bioscience) regenerating system containing NADH (0.2 mM, Sigma-Aldrich), PEP (0.2 mM, Sigma-Aldrich) and PK/LDH (6 U/mL / 10 U/ml, Carl Roth). The measurement was carried out at 37°C and a wavelength of 340 nm. The slit was chosen to be 1 nm and measurement intervals were set to 1 second. As NADH shows strong absorption at 340 nm, its consumption (and in an indirect manner, ATP’s consumption as well) can be monitored.

1 μM of Hsp902 (i.e. 2 μM monomers) or Cdc37_2_-Hsp90_2_-fusion were used with 1 μM or equimolar concentrations of the respective binding partners. After 20 minutes, radicicol (200 μM, Sigma) was added to inhibit Hsp90’s ATPase activity. The inhibition curve was used as reference and was subtracted from the data.

A one-way ANOVA was performed to compare the effect of nine different protein conditions on Hsp90’s ATPase rate. The one-way ANOVA revealed that there was a statistically significant difference in mean ATPase between at least two conditions (F(8,58) = [23.41464], p = 0.05). Tukey’s HSD Test for multiple comparisons found the conditions whose mean ATPase rate were significantly different at a significance level of p = 0.01, p = 0.05 and p = 0.1, respectively. All conditions shown in Fig. S1, selected ones in Fig. 4.

### ATPase assays to check protein fusions’ functionality

To check whether the introduction of the artificial linker alters the proteins’ interactions, ATPase assays of Hsp90_2_ only, Cdc37_2_-Hsp90_2_-fusion, and Hsp90_2_ in presence of freely added Cdc37 were compared both in absence and presence of the client kinase Ste11 (Fig. S1).

The addition of equimolar Cdc37 significantly reduces Hsp90’s already low rate of 1.3 ± 0.3 ATP/minute to 0.7 ± 0.3 ATP/minute. The linked Cdc37_2_-Hsp90_2_-fusion protein yields the very same result, clearly indicating that it is functional and that the protein interactions are not altered by the linker. The addition of equimolar Ste11 to Hsp90 also reduces its ATPase activity to 1.0 ± 0.3 ATP/min, although not significantly. However, when both Ste11 and Cdc37 are present, Cdc37’s reducing effect on Hsp90’s ATPase activity is abrogated. This holds true for the freely added Cdc37 as well as for the fusion-construct with rates of 1.1 ± 0.4 and 0.9 ± 0.3 ATP/min, respectively, indicating that Ste11 influences the interaction (and possibly also manner and/or position of binding) between Hsp90 and its cochaperone.

