## Supplementary Figure S1 for "Cochaperones convey the energy of ATP hydrolysis for directional action of Hsp90"

**Supplementary Information**

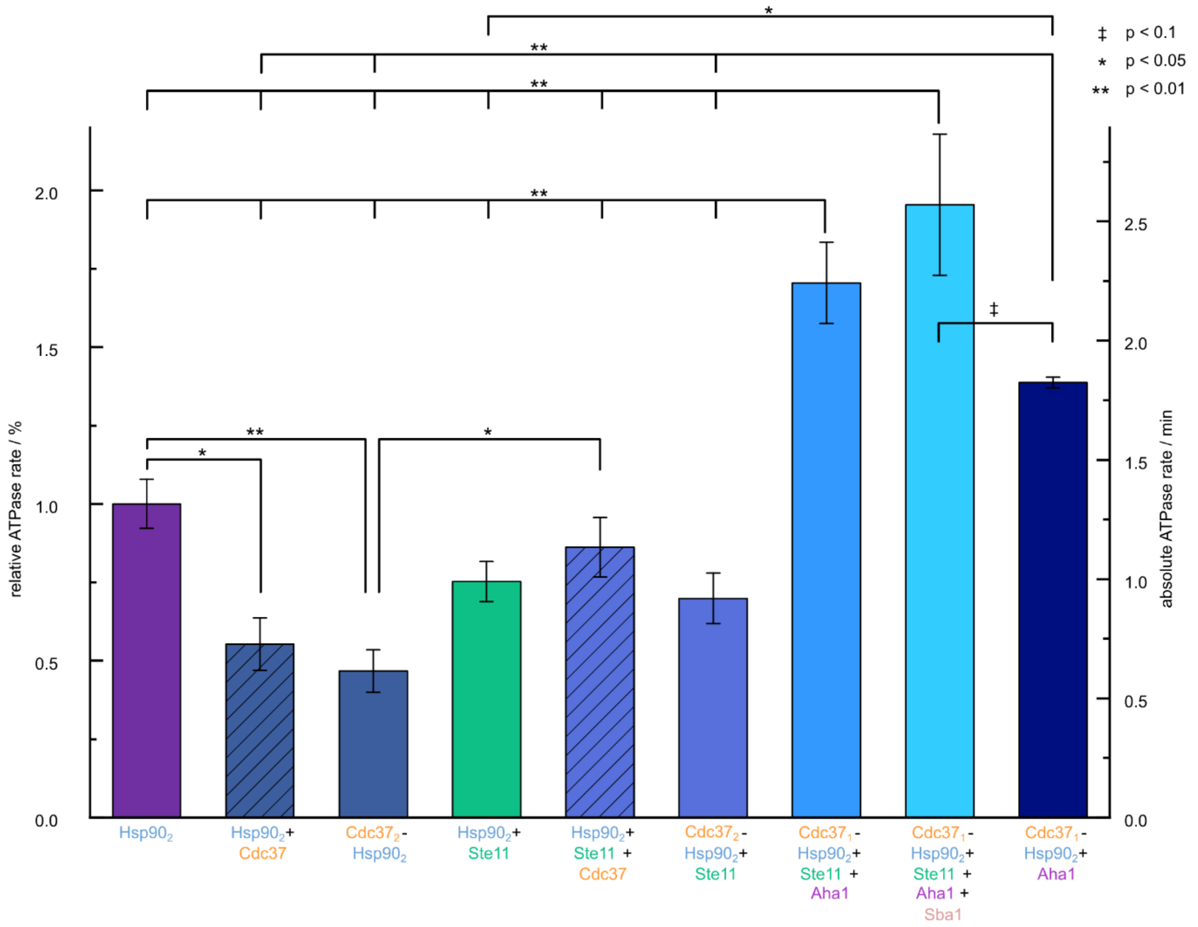


**Figure S1**: Relative and absolute ATPase rates of Hsp90 (2 µM) in presence and absence of cochaperones Cdc37, Aha1 and Sba1 and the client kinase Ste11.

$Hsp90_2$ (1 µM, i.e. 2 µM monomers) is a slow ATPase with 1.3 hydrolysed ATP per minute (violet). The addition of free Cdc37 (2 µM, equimolar) further decreases the ATPase rate (dark blue, striped). As the $Cdc37_2$-$Hsp90_2$-fusion shows the same behaviour, the functionality of this construct (dark blue) is proven. Ste11 (2 µM) slightly decreases Hsp90's ATPase activity (green). Additionally, Ste11 hinders Cdc37's ATPase-decreasing effect on Hsp90 of both the freely added protein (middle blue, striped) as well as the $Cdc37_2$-$Hsp90_2$-fusion (middle blue). Aha1, Cdc37, Sba1 and Ste11 together (1 µM each, lightest blue) strongly increase the ATPase to 2.6 ATP/min. This effect is not achieved by Cdc37 and Aha1 alone (1 µM each, darkest blue). Statistical significance tested by one-way ANOVA and Tukey post hoc test (described in methods).
